# A general method to develop excitation ratiometric neuropeptide sensors with enhanced sensitivity for *in vivo* applications

**DOI:** 10.64898/2026.09.17.752296

**Authors:** Huan Wang, Yuqi Yan, Yulin Zhao, Di Wang, Changhao Huang, Yulong Li

## Abstract

Neuropeptides regulate a wide range of physiological processes throughout the nervous and endocrine systems, yet their spatiotemporal dynamics *in vivo* remain poorly understood. While recently developed intensiometric sensors can detect neuropeptides, their use *in vivo* is often confounded by artifacts induced by hemodynamic changes, pH fluctuations, and motion. Here, we present a general strategy for engineering dual-excitation ratiometric (Ex-ratiometric) neuropeptide sensors by tuning the excited-state proton transfer properties of the fluorescent reporter. Dual excitation at 405 nm and 488 nm produces a ratiometric signal that intrinsically corrects fluorescence fluctuations unrelated to ligand binding. Using an AlphaFold3-guided in silico design approach combined with experimental validation, we developed a suite of Ex-ratiometric sensors. As a representative example, Ex-NTS2.0 enables robust detection of neurotensin (NTS) *in vivo* while remaining largely resistant to hemodynamics, pH, and motion-induced artifacts. Overall, these findings establish a scalable and mechanistically grounded platform for developing new Ex-ratiometric tools and provide a broadly applicable strategy for sensitive detection of neuropeptide dynamics in living systems.

## INTRODUCTION

Neuropeptides are a diverse class of signaling molecules that act as critical regulators of both the nervous and endocrine systems^1,2^. In the brain, neuropeptides modulate a broad range of physiological processes, from homeostatic processes such as feeding and metabolism to complex behaviors including sleep, reproduction, and cognitive processing^3,4^. Given their fundamental role in neuromodulation, neuropeptide signaling, primarily mediated by G protein-coupled receptors (GPCRs), represents a major therapeutic target for disorders such as chronic pain, obesity, and neurodegenerative diseases^5–10^.

Despite their importance, the precise spatiotemporal dynamics of neuropeptide release *in vivo* remain limited. Traditional detection methods such as microdialysis lack sufficient temporal and spatial resolution, limiting their ability to accurately capture rapid, localized signaling events^11,12^. Recently developed genetically encoded neuropeptide sensors have begun to address this challenge by enabling real-time monitoring of neuropeptide release with high spatiotemporal resolution^13–17^. However, most existing sensors rely on single-wavelength intensiometric readouts, making them particularly susceptible to artifacts in vivo. This limitation is particularly pronounced for neuropeptides, whose low endogenous concentrations often produce subtle fluorescence changes that are easily masked by hemodynamic fluctuations, pH changes, and motion artifacts^18–21^.

Hemodynamic changes often associated with changes in brain states, including sleep, wakefulness, and anesthesia^22–24^, alter tissue absorption and scattering properties, leading to non-specific changes in fluorescence intensity^25–27^. Neural activity also induces compartment-specific intracellular pH shifts with acidification occurring in soma^28–31^, and transient alkalinization observed at synaptic terminals^32^. These pH changes can directly affect the fluorescence of GFP or one of its variants-based sensors, whose chromophores are intrinsically sensitive to protonation state^33–35^. For instance, enhanced green fluorescent protein (EGFP) has a p*K*_a_ of approximately 6.0^34^; consequently, neuronal activity–induced intracellular acidification can trigger a non-specific decrease in its fluorescence intensity^36^. Furthermore, animal movement can introduce substantial motion artifacts, while variability in sensor expression levels across various cell populations and/or individual animals further complicates interpretation of the absolute signal changes. Together, these factors can significantly confound the detection of neuropeptide release *in vivo*.

Ratiometric imaging provides a powerful strategy for mitigating these confound influences and achieving quantitative measurements^37–40^. Ratiometric sensors generally employ either emission-ratiometric or excitation-ratiometric designs. Emission-ratiometric sensors typically rely on the fusion of a responsive indicator with a reference fluorophore such as a red fluorescent protein (RFP)^39,40^. However, the inclusion of an additional fluorescent protein significantly increases the probe’s size and often introduces complications due to differences in physicochemical properties, such as maturation rates and pH sensitivity, between the two distinct fluorophores^41–44^. Excitation-ratiometric approaches offer a more streamlined alternative. The synthetic calcium indicator Fura-2 established the utility of excitation-ratiometric imaging by enabling quantitative measurements of intracellular calcium dynamics through dual-wavelength excitation at 340 nm and 380 nm^37^. For genetically-encoded sensors, a similar strategy can be implemented by exploiting the intrinsic excited-state proton transfer (ESPT) properties of wild-type GFP (wtGFP)^45^. Upon photoexcitation, the absorption of a photon triggers an ultra-fast redistribution of electron density, prompting the excited neutral chromophore to release a proton and enter its ionized form (Fig. 1A). The coexistence of neutral (protonated) and ionized (deprotonated) chromophore states, each characterized by distinct excitation spectra^46^, provides a basis for excitation-ratiometric measurements. As both excitation channels originate from the same fluorophore, they share the same spatial distribution, photochemical environment, and other common-mode artifacts, thereby improving measurement fidelity *in vivo*.

**Fig. 1.**
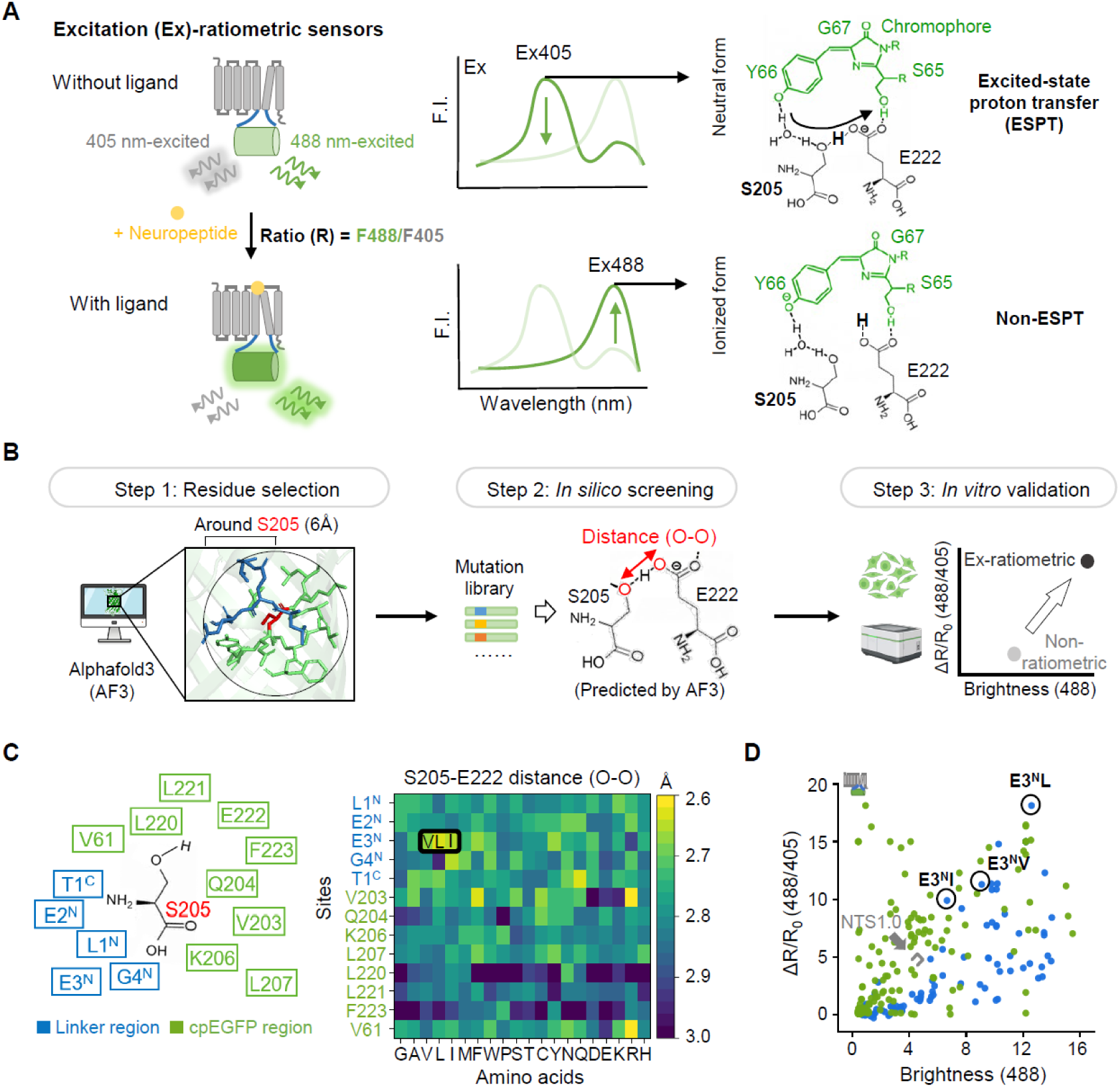
Computationally guided design of excitation-ratiometric neuropeptide sensors. A. Working principle and mechanism of Ex-ratiometric sensors. Left: schematic diagram depicting the dual-excitation strategy based on the GPCR scaffold in the ligand-free (i.e., unbound) and ligand-bound states, including the ratio formula (R = F488 / F405). Middle: excitation spectra curves of the sensor in the ligand-free state (top, dark green line) and ligand-bound (bottom, dark green line) states, indicating excitation wavelengths at 405 nm and 488 nm. Right: diagrams depicting the local hydrogen-bond networks in the neutral form (top) and ionized form (bottom) of wild-type GFP, showing residues Y66, S65, G67, S205, and E222, with the shared proton bolded. B. Three-step workflow for the *in silico* and *in vitro* screening of ESPT-modulating mutations using AlphaFold3 (AF3). Step 1: Selection of candidate residues within a 6-Å radius around residue S205. Step 2: *In silico* generation of the mutation matrix. Step 3: *In vitro* validation in HEK293T cells. C. *In silico* saturation mutagenesis screening corresponding to Step 2 in B. Left: structural map of selected candidate residues within a 6-Å radius around S205, with the linker and cpEGFP regions color-coded in blue and green, respectively. Right: heatmap of the predicted structural distances (O–O distance in Å) between the S205 and E222 side-chain oxygen atoms across 20 amino acid substitutions at the 13 candidate sites. D. *In vitro* screening of the GRAB_NTS_ mutation library corresponding to Step 3 in B. Individual variants are plotted and color-coded by their mutation domains, with linker sites and cpEGFP sites shown in blue and green, respectively. The *y*-axis displays the maximum ratiometric response (ΔR/R_0_), and the *x*-axis displays the fluorescence brightness under 488-nm excitation.

Here, we present an AlphaFold3-assisted strategy combined with experimental validation to develop excitation-ratiometric neuropeptide sensors, by modulating the ESPT within cpEGFP-based reporters. Using this strategy, we developed Ex-NTS2.0, an excitation-ratiometric sensor for neurotensin (NTS) and demonstrated its ability to reliably monitor NTS dynamics under diverse physiological and pharmacological conditions, including isoflurane anesthesia, tail suspension, and cocaine exposure. To assess the generalizability of this approach, we further developed a series of ratiometric neuropeptide sensors, including sensors for detecting somatostatin (SST) and pituitary adenylate cyclase activating polypeptide (PACAP). Together, these results establish a broadly applicable platform for developing ratiometric GPCR-based sensors, and enable the precise and robust detection of neuropeptide dynamics *in vivo*.

## RESULTS

### Computationally assisted development of excitation-ratiometric sensors

Ex-ratiometric neuropeptide sensors exhibit two distinct excitation peaks, typically occurring near 405 nm and 488 nm, that undergo opposing fluorescence changes upon ligand binding (Fig. 1A). In contrast, most genetically encoded neuropeptide sensors based on cpEGFP are intensiometric, showing a fluorescence increase only at 488 nm upon ligand binding^13,14^. This lack of a ratiometric response measured at 405 nm and 488 nm occurs because the chromophore in cpEGFP exists predominantly in the ionized form, leaving a limited number of chromophores in the neutral form and inherently suppressing the ESPT process. In GFPs, the chromophore exists in both a neutral form and an ionized form, with the neutral form maintaining a complex network of hydrogen bonds with surrounding residues such as S205 and E222 to facilitate ESPT^46,47^ (Fig. 1A). We therefore hypothesized that targeted modulation of this network might support the rational design of Ex-ratiometric sensors.

To identify target residues for modulating the ESPT, we established a three-step pipeline integrating structural prediction and experimental high-throughput screening (Fig. 1B). First, we used AlphaFold3 to predict the structure of the cpEGFP domain in GRAB (GPCR activation–based) sensors. Because the proton transfer requires precise hydrogen-bond positioning (typically within 3 Å), we screened all residues within a 6-Å radius of S205 (excluding E222) to identify candidate residues that might affect proton transfer. This approach identified 13 candidate amino acids, which were further categorized as residing in the linker or cpEGFP region (Fig. 1C). Next, we used *in silico* saturation mutagenesis to evaluate all possible amino acid substitutions at these 13 sites. We prioritized mutations predicted to decrease the distance between the side chain oxygen atoms of residues S205 and E222, which are critical for hydrogen-bond formation and ESPT efficiency. Finally, the candidate mutations were validated experimentally using saturation mutagenesis and high-content imaging by expressing the sensors in HEK293T cells.

AlphaFold3-based structural analysis revealed 13 candidate sites within 6-Å radius surrounding S205 except for the E222. During *in silico* screening, mutations at these 13 candidate amino acids indeed showed varied O–O distances. Notably, substitution of the glutamic acid—the third residue in the N-terminal linker (E3^N^)—with a hydrophobic amino acid such as leucine (L) or isoleucine (I) reduced the predicted O–O distance to 2.6–3.0 Å (Fig. 1C), a distance favorable for hydrogen bonding and proton transfer. Using the GRAB_NTS_ sensor as a prototype, large-scale screening identified E3^N^L as the optimal variant (Fig. 1D) which significantly increased the sensor’s Ex-ratiometric property, increasing the fluorescence response to 405-nm excitation by nearly 200% and increasing the ratiometric response by 300% compared to the NTS1.0 sensor (Fig. 1D). Importantly, the fluorescence response induced by excitation at 405 nm in the E3^N^ variants was strongly correlated with the predicted O–O distance (Fig. S1A), validating this AlphaFold3-assisted excitation-ratiometric sensor engineering approach.

### Characterization of the GRAB_Ex-NTS2.0_ sensor in cultured cells

When expressed in HEK293T cells, the Ex-NTS2.0 sensor localized efficiently to the plasma membrane and exhibited robust ratiometric properties (Fig. 2A). The excitation spectrum displayed two prominent peaks near 400 nm and 500 nm; upon ligand binding, the fluorescence intensity induced by 405-nm excitation decreased, while the intensity induced by 488-nm excitation increased (Fig. 2B). Compared to the NTS1.0 sensor, Ex-NTS2.0 had a substantially larger response in both channels, with maximum changes of +1200% at 488 nm and -35% at 405 nm, resulting in a 2000% ratiometric response with an apparent ligand affinity of 20 nM (Fig. 2C). Furthermore, the signal-to-noise ratio (SNR) of Ex-NTS2.0 was significantly higher compared to the SNR of NTS1.0 (Fig. 2D). We also generated a ligand-insensitive version, NTSmut, by introducing the Y342^7.13^R substitution in Ex-NTS2.0 (Fig. S1B), which serves as a control to verify the specificity of the signals observed *in vitro* and *in vivo*; we confirmed that NTSmut localizes to the plasma membrane but produces no detectable response to NTS even at concentrations up to 1 µM (Fig. S1C). When expressed in cultured cortical neurons, Ex-NTS2.0 traffics to the plasma membrane and produces a response profile and SNR comparable to those observed when expressed in HEK293T cells (Fig. S2).

**Fig. 2.**
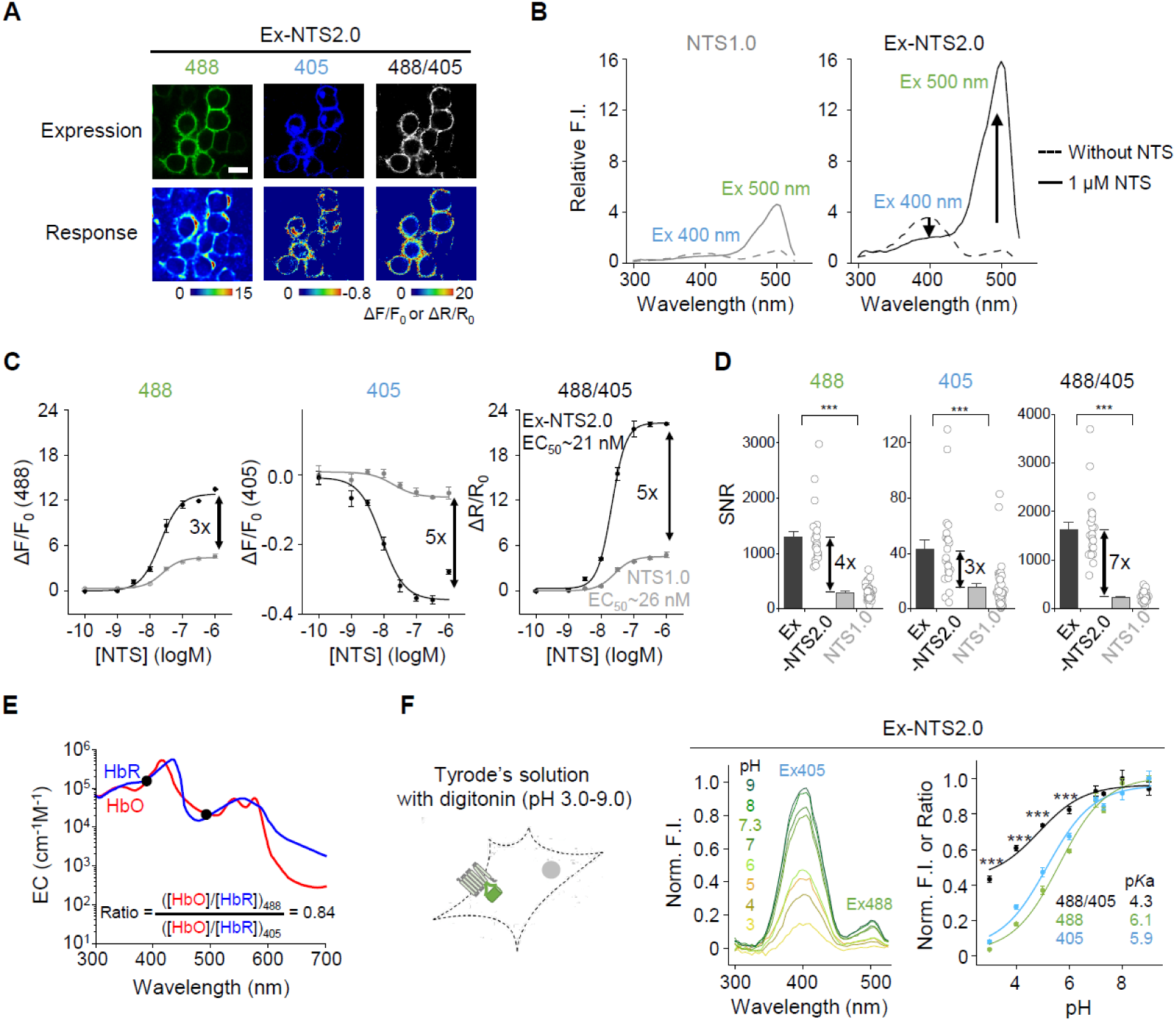
Ex-NTS2.0 exhibits high ratiometric responses and stability *in vitro*. A. Fluorescence and ratiometric images of Ex-NTS2.0 expressed in cultured HEK293T cells. Top: representative fluorescence images under 488-nm excitation (green) and 405-nm excitation (blue), along with the calculated grayscale ratio map (F488/F405). Bottom: pseudocolor maps (ΔF/F_0_ or ΔR/R_0_) showing the response after application of 1 μM NTS. Scale bar, 20 μm. B. Excitation spectral profiles of Ex-NTS2.0 (black lines) and NTS1.0 (gray lines) expressed in HEK293T cells. Dashed lines: baseline measurements in the absence of ligand; solid lines: measurements in the presence of 1 μM NTS. C. Dose-response curves of Ex-NTS2.0 (black curves) and NTS1.0 (gray curves) expressed in HEK293T cells under increasing concentrations of NTS. From left to right: the single-wavelength fluorescence changes (ΔF/F_0_) under 488-nm and 405-nm excitation, and the calculated ratiometric change (ΔR/R_0_), plotted against NTS concentration; n = 3 wells per group, 200–500 cells per well. D. Summary of the signal-to-noise ratio (SNR) measured for Ex-NTS2.0 (black bars) and NTS1.0 (gray bars) expressed in HEK293T cells. From left to right: the calculated SNR values obtained under 488-nm and 405-nm excitation, and the 488/405 ratiometric channel; n = 3 wells per group, 200–500 cells per well. \*\*\**P*<0.001 (two-tailed Student’s *t*-test). E. Absorption spectra and derived coefficient ratios for oxy-hemoglobin (HbO, red) and deoxy-hemoglobin (HbR, blue). Left vertical axis plots molar extinction coefficient (EC) on a logarithmic scale. F. Characterization of pH sensitivity measured for the Ex-NTS2.0 sensor expressed in HEK293T cells. Left: schematic diagram depicting cell permeabilization in digitonin-containing Tyrode’s solution (pH 3.0–9.0). Middle: excitation spectral profiles measured in the indicated pH values. Right: normalized titration curves versus pH for the 488-nm excitation channel (Ex488, green), the 405-nm excitation channel (Ex405, blue), and the calculated ratio channel (Ratio, black); n = 3 wells per group, 200–500 cells per well. \*\*\**P*<0.001 (one-way repeated measures ANOVA followed by Tukey’s multiple-comparison tests).

Pharmacologically, Ex-NTS2.0 retains the high ligand specificity of its parent receptor, NTS1R, showing negligible activation by other neuropeptides (Fig. S3). To investigate whether the Ex-NTS2.0 sensor couples to downstream signaling pathways, we used the luciferase complementation assay^48^ and the Tango assay^49^ to measure activation of the GPCR-mediated Gq and β-arrestin pathways, respectively. In contrast to the wild-type NTS1R receptor, which exhibited robust dose-dependent downstream signaling, Ex-NTS2.0 has only negligible downstream coupling (Fig. S4). Together, these results demonstrate that Ex-NTS2.0 is a highly sensitive, ligand-specific tool for detecting NTS dynamics, without causing measurable downstream signaling.

### Ex-NTS2.0 has high ratiometric performance and stability *in vitro*

As mentioned above, hemodynamic artifacts often pose a significant challenge for *in vivo* fluorescence imaging. Because hemoglobin exhibits strong wavelength-dependent absorption in the blue-green spectral range, fluctuations in blood volume and/or oxygenation can introduce substantial artifacts into fluorescence measurements^25,26^. Notably, the absorption spectra of oxyhemoglobin and deoxyhemoglobin are relatively balanced between the 405-nm and 488-nm channels (Fig. 2E). Consequently, the ratiometric readout produced by excitation at 488 and 405 nm effectively normalizes these intensity fluctuations, showing that Ex-NTS2.0 has the inherent ability to resist hemodynamic artifacts and maintain its signal sensitivity *in vivo*.

To evaluate the pH sensitivity of Ex-NTS2.0, we compared its properties with membrane-localized pH-sensitive (EGFP-CAAX) and pH-insensitive (Gamillus-CAAX) fluorescence proteins. Consistent with previous reports^50,51^, EGFP-CAAX has a p*K*_a_ of approximately 5.8, while Gamillus-CAAX has low pH sensitivity with a p*K*_a_ of approximately 3.1 (Fig. 3F). With respect to Ex-NTS2.0, we found that its individual excitation channels displayed pH sensitivity comparable to that of EGFP-CAAX. In contrast, the ratiometric signal exhibited significantly reduced pH sensitivity compared to either excitation channel alone, with a leftward shift in the pH-response curve and a p*K*_a_ of approximately 4.3 (Fig. 2F). Thus, Ex-NTS2.0 is highly stable under both pH changes and hemodynamic fluctuations.

**Fig. 3.**
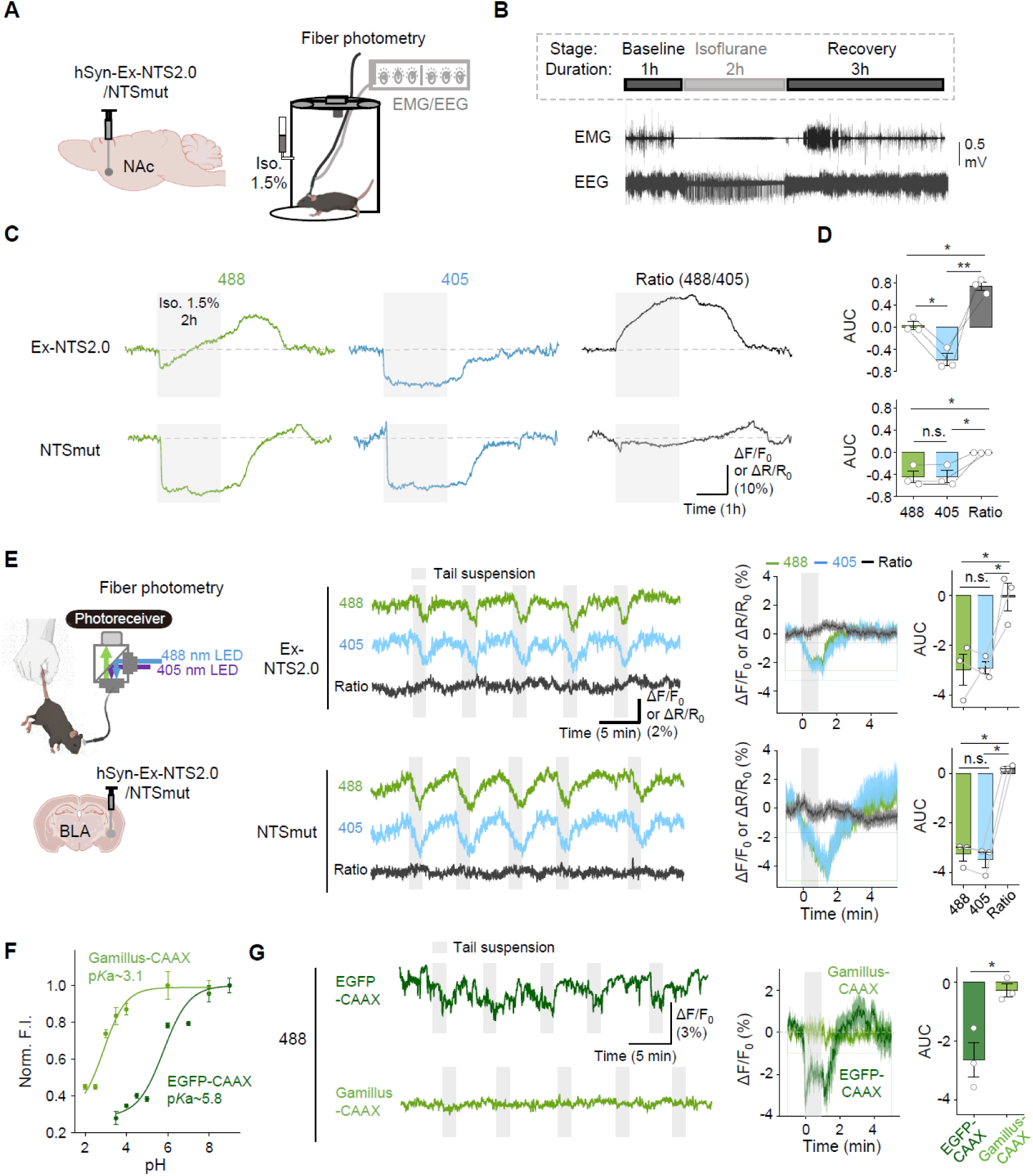
Ratiometric recording with Ex-NTS2.0 minimizes hemodynamic and pH artifacts. A. Experimental setup for *in vivo* fiber photometry and recording during anesthesia. Left: schematic diagram showing the viral injection of Ex-NTS2.0 or NTSmut into the NAc. Right: the recording chamber configuration with an isoflurane vaporizer (delivering 1.5% isoflurane) and an integrated EMG/EEG monitor. B. Timeline and representative traces of the anesthesia paradigm. Top: timeline showing the sequential stages and duration. Bottom: representative EMG and EEG data recorded throughout the experiment. C. Representative continuous fiber photometry traces (ΔF/F_0_ or ΔR/R_0_) recorded in the NAc in the 488-nm (green), 405-nm (blue), and ratio (black) channels for Ex-NTS2.0 (top) and NTSmut (bottom). Gray shaded blocks indicate the 2-h isoflurane window, and dashed lines indicate the baseline. D. Summary of the calculated AUC during anesthesia in the three indicated channels for Ex-NTS2.0 (top) and NTSmut (bottom). Individual animals are connected by gray lines. \**P*<0.05, \*\**P*<0.01, and n.s., not significant (one-way repeated measures ANOVA followed by Tukey’s multiple-comparison tests). E. Left: schematic diagram depicting the viral injection (Ex-NTS2.0 or NTSmut) into the BLA and the dual-excitation (488-nm and 405-nm) fiber photometry setup. Middle: representative continuous photometry traces (ΔF/F_0_ or ΔR/R_0_) for the 488 (green), 405 (blue), and ratio (black) channels across multiple trials in mice expressing Ex-NTS2.0 (top) or NTSmut (bottom). Gray shaded blocks indicate individual tail-suspension trials. Right: event-locked trial-averaged line traces and corresponding summary plots of the AUC quantified for the 488-nm, 405-nm, and ratio channels for Ex-NTS2.0 (top row) and NTSmut (bottom row). \**P*<0.05 and n.s., not significant (one-way repeated measures ANOVA followed by Tukey’s multiple-comparison tests). F. *In vitro* pH sensitivity of Gamillus-CAAX and EGFP-CAAX. Normalized fluorescence intensity (Norm. F.I.) plotted against pH values to determine the p*K*_a_ for Gamillus-CAAX and EGFP-CAAX. G. *In vivo* validation of activity-induced pH changes during tail-suspension stress. Left: Representative in vivo fiber photometry traces (488 nm excitation channel) showing fluorescence changes (ΔF/F_0_) of EGFP-CAAX (top, dark green) and Gamillus-CAAX (bottom, light green) expressed in the BLA. Middle: Averaged fluorescence responses aligned to the onset of the tail-suspension stimulus (shaded gray bar). Right: Group data quantifying the AUC for EGFP-CAAX and Gamillus-CAAX during the stress paradigm. Individual circles represent single animals. \**P*<0.05 (two-tailed Student’s *t*-test). Data are shown as mean±SEM.

### Ratiometric recording with Ex-NTS2.0 is unaffected by hemodynamic and pH artifacts *in vivo*

While the ratiometric strategy serves to suppress the effects of environmental noise, its dual-excitation nature may also enable the sensor to amplify true biological signals against the background of artifacts induced by physiological fluctuations. To investigate this possibility, we examined whether Ex-NTS2.0 can accurately monitor NTS dynamics under isoflurane anesthesia, a condition known to induce large changes in cerebral blood flow and hemoglobin concentration^23,52^. We therefore expressed Ex-NTS2.0 in the nucleus accumbens (NAc) and the ventral tegmental area (VTA) and continuously recorded the fluorescence signals during isoflurane induction and recovery (Fig. 3A-B, Fig. S5). When expressed in the NAc, Ex-NTS2.0 exhibited opposing fluorescence changes in the two excitation channels, resulting in an increased ratiometric response (Fig. 3C). Moreover, quantification of the area under the curve (AUC) revealed that the change in the ratio was significantly larger than either excitation channel alone (Fig. 3D).

Notably, despite isoflurane-induced hemodynamic artifacts, which were prominent as non-specific decreases in both channels in the NTSmut control sensor expressed in the NAc, the ratiometric signal was remarkably stable (Fig. 3C). Interestingly, in the VTA we observed large fluctuations in the single channels, but no change in the ratio, indicating that the intramolecular self-calibration mechanism of Ex-NTS2.0 effectively minimized these anesthesia-induced artifacts (Fig. S5A-B). Group analyses further revealed that the AUC of the Ex-NTS2.0 signal in the NAc was significantly larger than that of both the NTSmut control and the Ex-NTS2.0 signal recorded in the VTA, confirming brain region–specific NTS release induced by isoflurane (Fig. S5C). These results demonstrate that Ex-NTS2.0 can reliably monitor NTS release in the NAc while effectively canceling out anesthesia-induced hemodynamic artifacts via its ratiometric self-calibration mechanism.

We next assessed whether Ex-NTS2.0 is vulnerable to pH-induced artifacts *in vivo* using a tail-suspension stress paradigm, as acute stress is known to trigger robust neuronal activation and related intracellular acidification in the basolateral amygdala (BLA). Upon expressing Ex-NTS2.0 in the BLA, we found that although both excitation channels exhibited stimulus-evoked decrease in signal intensity, the ratiometric signal remained remarkably stable across repeated trials; similar results were obtained when we expressed the control sensor, NTSmut (Fig. 3E). To validate these findings against fluorescent proteins, we performed control experiments using the pH-sensitive EGFP-CAAX and pH-insensitive Gamillus-CAAX. EGFP-CAAX showed a significant decline in fluorescence in the BLA following each stimulus, while the Gamillus-CAAX signal was unchanged (Fig. 3G), confirming the presence of an activity-induced pH drop. These results demonstrate that Ex-NTS2.0’s ratiometric measurement is unaffected by changes in pH, improving the fidelity of the neuropeptide signal measured *in vivo*.

### Ex-NTS2.0 can be used to monitor cocaine-induced NTS release *in vivo*

Because NTS signaling within the reward circuitry has been implicated in drug addiction and associated behavioral changes^53,54^, we investigated whether Ex-NTS2.0 can be used to accurately and stably monitor NTS dynamics under conditions of increased locomotion. We first expressed Ex-NTS2.0 in the NAc and then recorded the sensor using fiber photometry in freely behaving mice (Fig. 4A-B). We found that repeated cocaine administration induced a significant increase in locomotor activity, accompanied by a ratiometric signal. Specifically, NTS release triggered opposing fluorescence changes in the sensor’s two channels, with an increased signal under 488-nm excitation and a decreased signal under 405-nm excitation. The opposing responses between the two channels increased the ratiometric dynamic range, improving signal fidelity under conditions of substantial motion (Fig. 4C). Notably, we observed a progressive increase in the AUC of the fluorescence response across the 488-nm channel, the 405-nm channel, and the ratio between channels measured between the first and second cocaine injections, suggesting a progressive potentiation of NTS release upon each subsequent pharmacological challenge (Fig. 4C-E, Fig. S6A).

**Fig. 4.**
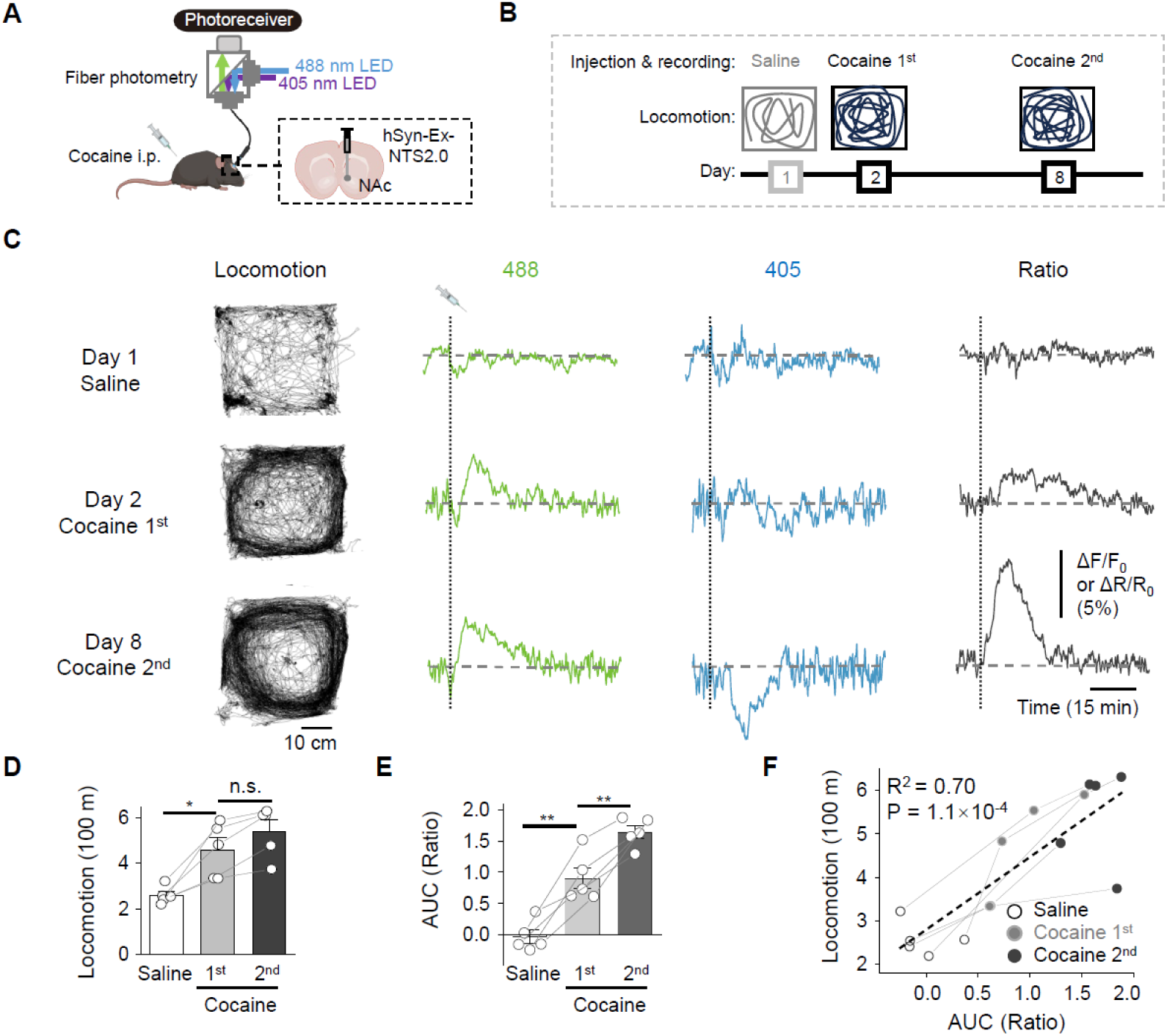
Ex-NTS2.0 monitors robust cocaine-induced NTS release *in vivo*. A. Schematic diagram depicting *in vivo* fiber photometry recording in the NAc before and after intraperitoneal injections of saline of cocaine, showing the dual-excitation (488-nm and 405-nm) optical configuration. B. Experimental timeline showing sequential open-field locomotion tracking and optical recording on day 1 (saline), day 2 (1^st^ cocaine injection), and day 8 (2^nd^ cocaine injection). C. Behavioral tracking maps and representative simultaneous traces (ΔF/F_0_ or ΔR/R_0_) in response to following saline and cocaine injections. From left to right: locomotor trajectories, the 488-nm channel (green), the 405-nm channel (blue), and the ratio channel (black). D-E. Summary of total locomotor distance and calculated AUC values in the ratio channel following the saline and cocaine injections. Individual animal points are connected by gray lines. \**P*<0.05, \*\**P*<0.01, and n.s., not significant (one-way repeated measures ANOVA followed by Tukey’s multiple-comparison tests). F. Correlation plots between locomotor distance traveled and calculated AUC values in the ratio channel.

Quantitative analyses revealed a significant correlation between locomotor activity and the AUC of the fluorescence signals for both excitation channels and the ratio between channels (Fig. 4F, Fig. S6B), demonstrating that Ex-NTS2.0 can maintain high signal fidelity even during intense motion. To confirm the specificity of these signals, we expressed the NTSmut control sensor in the NAc and found that neither saline nor cocaine administration elicited any significant changes in NTSmut fluorescence (Fig. S7), confirming that the signals observed with Ex-NTS2.0 are specifically driven by endogenous NTS release rather than motion-related or non-specific optical artifacts. Together, these results demonstrate that Ex-NTS2.0 is a self-correcting tool that can be used to reliably and sensitively detect NTS-associated signals under complex *in vivo* conditions.

### Generalization of the design strategy to develop additional neuropeptide sensors

To evaluate the generalizability of this approach, we applied site-saturated mutagenesis to the E3^N^ site in six additional neuropeptide sensors for detecting somatostatin (SST), neuropeptide Y (NPY), pituitary adenylate cyclase activating polypeptide (PACAP), vasoactive intestinal peptide (VIP), cholecystokinin (CCK) and corticotropin releasing factor (CRF) (Fig. 5A). Consistent with our findings for Ex-NTS2.0, structural predictions indicated that hydrophobic substitutions—particularly leucine and isoleucine—at the E3^N^ site reduced the distance between the side-chain oxygen atoms in S205 and E222 for all sensors tested (Fig. 5A). Experimental screening confirmed that these E3^N^ variants had a broadly increased ratiometric dynamic range for all neuropeptide sensors tested, especially for the variants with leucine or isoleucine (Fig. 5B). As a representative example, we expressed the Ex-SST2.0 sensor in HEK293T cells and measured opposing fluorescence changes under 405-nm and 488-nm excitation upon SST application (Fig. 5C-D). This SST sensor achieved a maximum ratiometric response of 1600%, with an apparent affinity of approximately 152 nM (Fig. 5E). Furthermore, the ratiometric signal produced by Ex-SST2.0 had significantly higher pH stability compared to the sensor’s single-channel measurements, with a p*K*_a_ of approximately 3.4 (Fig. 5F). These results therefore identify E3^N^ site as a critical site for tuning the ratiometric properties of dual-excitation neuropeptide sensors.

**Fig. 5.**
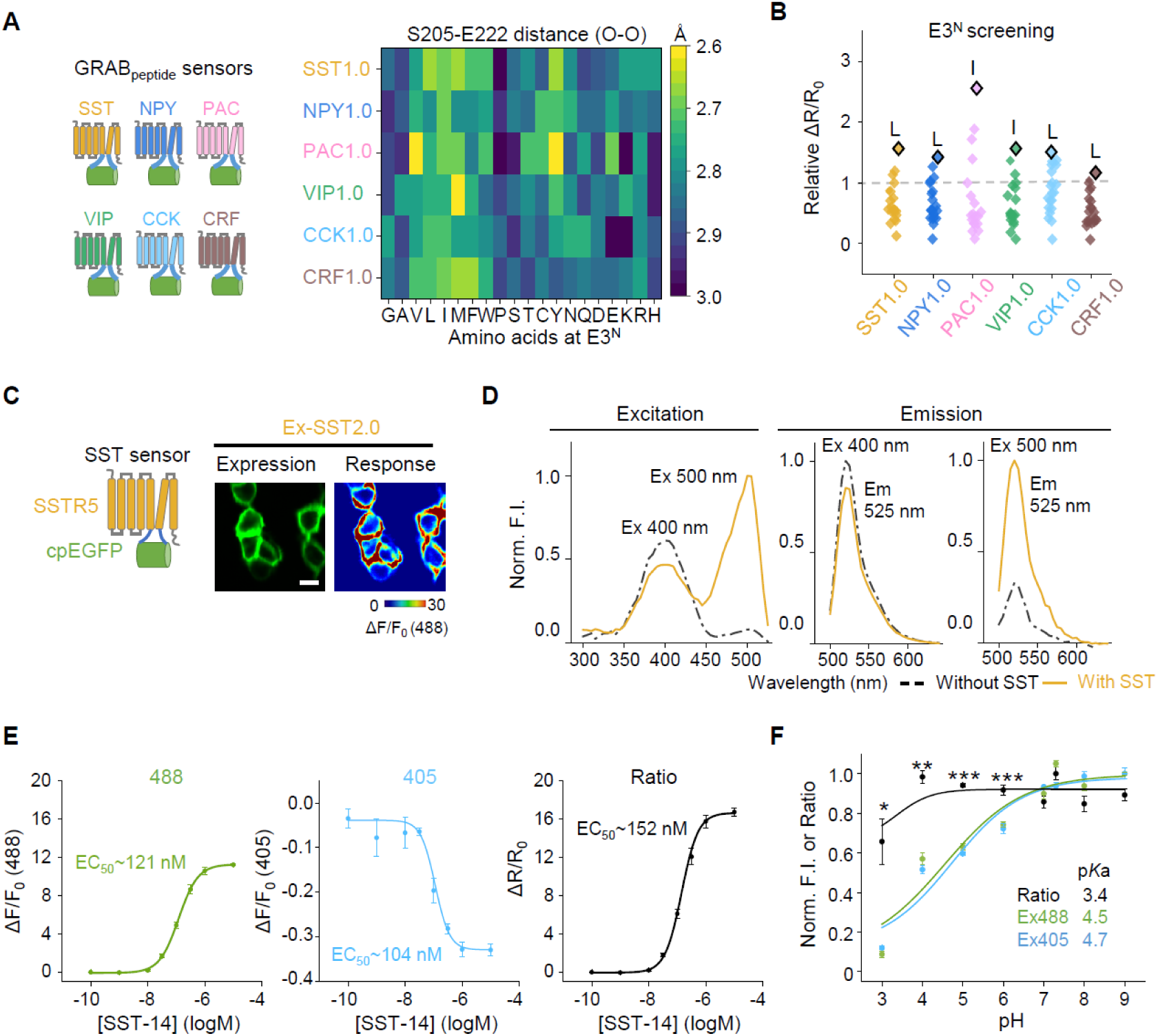
Generalization of the design strategy to diverse neuropeptide sensors. A. *In silico* saturation mutagenesis screening matrix at the E3^N^ site for the indicated six neuropeptide sensors. Left: schematic diagrams of the tested GPCR scaffolds. Right: heatmap showing AF3-predicted structural O–O distances between the S205 and E222 side-chain oxygen atoms for 20 amino acid substitutions at the E3^N^ position in the individual sensor scaffolds. B. Screening validation showing the relative ratiometric dynamic ranges (relative ΔR/R_0_) of individual amino acid mutations at the E3^N^ site for all six sensors. C. Expression and response images of the Ex-SST2.0 sensor expressed in HEK293T cells. Left: structural diagram based on the SSTR5 scaffold. Middle: representative baseline fluorescence image showing cell-surface trafficking. Right: calculated ΔF/F_0_ (488-nm) fluorescence response map after application of 1 μM SST-14. Scale bar, 20 μm. D. Excitation (left) and emission (right) spectral profiles of Ex-SST2.0 measured in the absence of ligand (black dashed lines) and in the presence of 1 μM SST-14 (yellow solid lines). E. Dose-response curves of Ex-SST2.0 measured under increasing concentrations of SST-14. From left to right: the fluorescence change (ΔF/F_0_) measured under 488-nm (green) and 405-nm (blue) excitation, and the calculated ratiometric change (ΔR/R_0_, black); n = 3 wells per group, 200–500 cells per well. F. Normalized titration curves plotted against pH for Ex-SST2.0 in the 488-nm channel (green), the 405-nm channel (blue), and the calculated ratio channel (black); n = 3 wells per group, 200–500 cells per well. \**P*<0.05, \*\**P*<0.01, and \*\*\**P*<0.001.

In summary, we present a generalizable strategy for engineering excitation-ratiometric neuropeptide sensors by targeting E3^N^ as a key structural hotspot. The robust performance of Ex-NTS2.0 *in vivo* confirms that this sensor can be used to provide artifact-resistant, high-fidelity monitoring of neuropeptide dynamics despite changes in hemodynamics, pH, and motion. Finally, our AlphaFold3-assisted approach was successfully used to generate six additional neuropeptide sensors, yielding a significantly larger ratiometric dynamic range and improved pH stability exemplified by the Ex-SST2.0 sensor.

## DISCUSSION

Here, we introduced a mechanistically driven strategy for converting intensiometric neuropeptide sensors into excitation-ratiometric sensors by modulating the ESPT property of cpEGFP. By tuning the local hydrogen-bond network surrounding the chromophore, we generated a bidirectional fluorescence response under dual excitation at 488 nm and 405 nm, providing an internal self-calibration mechanism that is particularly advantageous for *in vivo* applications.

Single-wavelength intensiometric sensors are highly prone to non-specific optical artifacts *in vivo* originating from hemodynamic fluctuations, pH alterations and animal motion. Given the picomolar-to-nanomolar endogenous concentrations of neuropeptides^18–21^, their modest specific fluorescence changes are easily overwhelmed by these non-specific interferences. Our results demonstrate that our Ex-ratiometric sensors can effectively overcome these challenges by enabling intrinsic normalization. Because the two excitation channels respond in opposite directions to ligand binding—but respond in the same direction to most forms of common-mode noise—ratiometric processing effectively cancels out non-specific artifacts such as hemodynamic fluctuations and tissue movement. While retaining the high affinity and specificity of the previous intensiometric sensor NTS1.0, Ex-NTS2.0 significantly increases the signal-to-noise ratio and provides a robust solution for high-fidelity, quantitative monitoring of neurochemical dynamics under complex *in vivo* conditions.

In addition to their self-correcting capability, the Ex-ratiometric sensors have several distinct advantages. First, compared to other ratiometric sensors such as FRET-based sensors, which often suffer from a limited dynamic range and complex spectral overlap^55,56^, our Ex-ratiometric approach maintains a high dynamic range (ΔR/R_0_) within a streamlined single fluorescent protein scaffold. Second, the dual-excitation spectra at 488 nm and 405 nm is compatible with widely available imaging modalities, including wide-field microscopy and fiber photometry systems^55,57,58^, increasing the sensor’s experimental applications without the need for specialized optical configurations. Furthermore, given the conserved structural dynamics of ligand-induced GPCR activation and the systematic principles underlying our engineering strategy^59,60^, this method can likely be extended to include the vast GPCR superfamily, including receptors for lipids, purines, and metabolic targets such as GLP-1, thus providing a comprehensive toolkit for precise neurochemical mapping.

Furthermore, the generalizable framework of our excitation-ratiometric design—predicated on modulating the ESPT pathway within the cpEGFP scaffold—presents a universal paradigm that can be extended far beyond neuropeptide sensors. Extending this strategy to existing intensiometric indicators for classic neurotransmitters (e.g., glutamate, GABA) and monoamines (e.g., dopamine, serotonin) would yield highly reliable, self-calibrating tools capable of minimizing pH and hemodynamic artifacts *in vivo*. Crucially, this engineering principle will be also vertically applicable to intracellular sensors, such as those monitoring second messengers like cAMP or Ca^2+^. Collectively, this generalizable approach provides a systematic platform to convert existing intensiometric sensors into next-generation, artifact-resistant ratiometric tools.

By expressing Ex-NTS2.0 *in vivo*, we monitored NTS dynamics with high sensitivity and accuracy. Our findings demonstrate the importance of using ratiometric measurements by separating true biological signals from imaging artifacts across different brain states. For instance, under isoflurane anesthesia the sensor successfully recorded bona fide NTS release dynamics within the NAc. In contrast, in the VTA we observed a clear dip in the raw signal; such a decrease could easily be misinterpreted as a ligand-induced decrease, but was shown by our Ex-ratiometric system to be caused by local hemodynamics-induced and/or pH-induced artifacts. This regional heterogeneity in NTS release likely reflects differences in neuronal origins and their projection patterns. In the NAc, NTS is primarily synthesized and released locally by D1 receptor–expressing medium spiny neurons; in contrast, in the VTA NTS depends more heavily on distal inputs from regions such as the lateral hypothalamus and the central nucleus of the amygdala^61,62^. Thus, isoflurane may selectively activate NTS-expressing neurons in the NAc or specifically facilitate their vesicular release. Given that the NAc is a critical hub not only for reward processing but also for sleep-wake transitions and anesthesia-induced state switching^63,64^, the cumulative release of NTS in this region may play a specialized role in maintaining the anesthetic state.

Although we show that our AlphaFold3-assisted engineering strategy is remarkably versatile with respect to increasing the ratiometric performance of a wide range of neuropeptide sensors, the precise biophysical mechanism by which the E3^N^ site modulates the ESPT property remains unclear. This uncertainty stems primarily from the fact that our current approach relies on static structural predictions rather than dynamic conformational analyses. Specifically, AlphaFold3 has inherent limitations in terms of modeling the chromophore’s mature state and explicitly incorporating essential water molecules^65^, both of which are essential for accurately predicting the complete proton-transfer network required for ESPT^46^. Consequently, future research integrating molecular dynamics simulations will be pivotal for advancing the rational design of new sensors. By capturing the transient and dynamic proton-transfer networks that are often obscured in static models, these integrated computational approaches will provide deeper insights at the atomic level and further refine the development of next-generation ratiometric tools.

Regarding these tools’ practical applications, although our Ex-ratiometric sensors can reliably and sensitively report neuropeptide dynamics *in vivo*, the 405-nm and 488-nm excitation channels still have inherent physical constraints under complex imaging conditions. On one hand, biological tissues produce significant autofluorescence in the blue-green spectrum^66^, which can weaken the signal-to-background ratio. On the other hand, the substantial scattering and absorption of short-wavelength light in dense tissues restrict the effective imaging depth^67,68^. In contrast, red-shifted fluorescent proteins offer superior spectral properties that are better suited for deep-tissue and multicolor imaging^69,70^. Consequently, future optimizations should include expanding the sensor toolkit’s spectral diversity, potentially through red-shifted Large Stokes Shift (LSS)-based sensors. Such advances would not only increase imaging depth but would also minimize 405-nm light-induced phototoxicity during long-term *in vivo* monitoring, thereby expanding the use of these sensors to include long-term studies of complex neurochemical signaling.

In conclusion, our Ex-ratiometric neuropeptide sensors provide a set of robust, artifact-resistant tools for reliably monitoring neuropeptide dynamics *in vivo*. By transitioning from an intensiometric measure to a ratiometric approach, we introduce a series of high-fidelity tools for dissecting the complex spatiotemporal dynamics of neuropeptides in both health and disease.

## METHODS

### Mice

All experimental procedures involving animals were conducted in accordance with protocols approved by the Animal Care and Use Committee of Peking University. Wild-type C57BL/6J mice and newborn Sprague-Dawley rats (P0) were purchased from Beijing Vital River Laboratory Animal Technology Co., Ltd. Mice were used for the *in vivo* experiments, while P0 rat pups were used to prepare the primary cortical neuron cultures. All animals were housed in a temperature-controlled (18–23°C) and humidity-controlled (40–60%) facility under a 12-h light/dark cycle, with *ad libitum* access to food and water.

### Cell lines

HEK293T cells (CRL-3216, ATCC) were used to generate stable cell lines expressing various neuropeptide sensors. These stable lines were created using the PiggyBac transposon system. In brief, cells were co-transfected with a hyperactive piggyBac transposase (pCS7-PiggyBAC, containing the S103P and S509G mutations) and a donor vector containing the sensor’s coding sequence driven by the CAG promoter, followed by an IRES-puromycin selection cassette, all flanked by PiggyBac inverted terminal repeat (ITR) sequences. Stable transformants were selected using 1 μg/ml puromycin starting 24 h post-transfection. The HTLA cell line, which was for the Tango assay, was a generous gift from Bryan L. Roth. All cell lines were maintained in DMEM (Biological Industries) supplemented with 10% (v/v) fetal bovine serum (FBS; CellMax) and 1% (v/v) penicillin-streptomycin (Gibco). Cells were cultured at 37℃ in humidified air containing 5% CO_2_.

### Cultured rat primary cortical neurons

Primary cortical neurons were prepared from P0 Sprague–Dawley rat pups of both sexes (Beijing Vital River Laboratory Animal Technology Co., Ltd.). In brief, the brains were removed and the cortices were carefully dissected in ice-cold phosphate-buffered saline (PBS). The tissue was dissociated using 0.25% trypsin-EDTA (Gibco), and the resulting cell suspension was plated onto glass coverslips pre-coated with poly-D-lysine hydrobromide (Sigma). The neurons were cultured in Neurobasal medium (Gibco) supplemented with 2% B-27, 1% GlutaMAX, and 1% penicillin-streptomycin (all from Gibco). The neurons were cultured at 37℃ in humidified air containing 5% CO_2_.

### AAV expression

Recombinant adeno-associated viruses (AAV, serotype 9) expressing the neuropeptide sensors were packaged and purified by WZ Biosciences with genomic titers ranging from 3-5 x 10^13^ v.g./ml. For expression in primary rat cortical neurons, the viral vectors were added to the culture medium at DIV 5–7; the neurons were then cultured for an additional 7–10 days post-infection to allow for robust sensor expression and proper membrane trafficking prior to imaging experiments.

### Molecular biology

The plasmids used in this study were synthesized primarily using the Gibson assembly method. DNA fragments were amplified via PCR using primers (Tsingke Biological Technology) designed with 25–30-bp overlapping sequences. These fragments were subsequently assembled using a reaction mixture of T5 exonuclease (New England Biolabs), Phusion DNA polymerase (Thermo Fisher Scientific), and Taq ligase (iCloning). All resulting constructs were verified using Sanger sequencing (Tsingke Biological Technology).

To evaluate the neuropeptide sensors expressed in HEK293T cells, cDNAs encoding the candidate sensors were cloned into the pDisplay vector, which features an upstream IgK leader sequence. A downstream IRES-mCherry-CAAX cassette was incorporated to anchor the reporter to the cell membrane and to provide a fluorescent signal for normalization of sensor intensity. For expression in cultured neurons, sequences for the lead neuropeptide sensors and their non-responsive variants were subcloned into the pAAV vector under the control of the human synapsin (*hSyn*) promoter.

For the Tango assay, genes encoding the wild-type neuropeptide receptors and their derived sensors were cloned into the pTango vector. For spectral characterization and the generation of stable cell lines, sensor sequences were cloned into the pPacific vector, which contains terminal repeats and an IRES-puromycin selection marker. Stable expression was achieved by co-transfection with a hyperactive piggyBac transposase (containing the S103P and S509G mutations in the pCS7-PiggyBAC vector; ViewSolid Biotech).

### Confocal imaging of cultured cells

HEK293T cells and primary rat cortical neurons were imaged using a Nikon confocal microscope (NIS-Element v4.51.00) equipped with a suite of objectives (20x air; 40x, 60x, and 100x oil). To capture the excitation-ratiometric signals, green fluorescence was sequentially or simultaneously excited using 488-nm and 405-nm lasers and collected via a 525/50-nm emission filter. Red fluorescence (mCherry) was excited using a 561-nm laser and collected via a 595/50-nm filter. Signals were detected using high-sensitivity photomultiplier tubes.

High-resolution images (1024 x 1024 pixels) were acquired before and after the application of saturating ligand concentrations using the 60x or 100x oil objective. Images were stabilized and aligned in ImageJ (NIH). Background-subtracted ratiometric images (R) were generated by calculating the pixel-by-pixel ratio of the 488-nm channel to the 405-nm channel (F488/F405). Response magnitude pseudo-color maps were calculated using ΔR/R_0_ = [(Rpost - Rpre) / Rpre] and displayed using the "Jet" look-up table (LUT).

Dynamic neuropeptide responses were imaged using a 20x air objective (512 x 512 pixels; 3-s interval) for 20–60 min. Following a stable baseline period, increasing concentrations of ligand were applied sequentially once the fluorescence reached a plateau. After background subtraction and channel splitting, fluorescence values (F) and ratios (R) were extracted from cell-specific regions of interest (ROIs). The sensor response was quantified as:

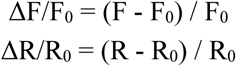

Dose-response curves were then generated by plotting the plateau response against the logarithmic ligand concentration.

### Spectra measurements

To characterize the neuropeptide sensors, HEK293T cells stably expressing the sensor under the control of the CAG promoter were seeded into 384-well plates in the presence or absence of 1 μM of the corresponding peptide ligand. For transient expression experiments, HEK293T cells were transfected with the sensor expression plasmid or an empty vector (as a background control) in 6-well plates. At 24–36 h post-transfection, the cells were dissociated using trypsin, washed with PBS, and resuspended in Tyrode’s solution with or without 1 μM peptide ligand before being transferred to 384-well plates. Fluorescence spectra were recorded using a Safire2 multi-mode plate reader (Tecan). Excitation spectra were acquired from 300 to 520 nm in 5-nm increments with a 20-nm bandwidth, while monitoring emission at 560 nm (20-nm bandwidth). Emission spectra were acquired from 500 to 700 nm in 5-nm increments (20-nm bandwidth) with the excitation wavelength set at either 390 nm or 455 nm (20-nm bandwidth) to capture the sensors’ ratiometric properties. Background fluorescence was determined from empty vector–transfected cells and was subtracted from all experimental groups. Tabulated molar extinction coefficient values (ε) of oxy- and deoxyhemoglobin were retrieved from a spectral dataset compiled by Scott Prahl (OMLC; https://omlc.org/spectra/hemoglobin/summary.html) and used to generate hemoglobin absorption spectra.

### MiniG protein luciferase complementation assay

To evaluate the coupling of neuropeptide sensors to downstream G proteins, a NanoBiT-based MiniG recruitment assay was performed. HEK293T cells were co-transfected with MiniG-LgBit and the SmBit-fused sensor or the wild-type GPCR. At 36–48 h post-transfection, the cells were harvested in PBS, supplemented with Furimazine substrate (Nano-Glo, Promega), and aliquoted into 96-well plates. Bioluminescence was measured using a Victor X5 reader (PerkinElmer, 1000-ms integration) following 10–15 min incubation with peptide ligands applied at 10 pM to 10 μM. Dose-response curves were fitted using a nonlinear function in OriginLab to determine the potency and efficacy of G protein recruitment for each neuropeptide sensor variant.

### Tango assay

To evaluate the coupling of neuropeptide sensors to downstream signaling pathways (e.g., beta-arrestin recruitment), Tango assays were performed. HTLA cells were transfected with plasmids encoding either the wild-type neuropeptide receptors or the corresponding sensors in 6-well plates. At 24 h post-transfection, the cells were dissociated using trypsin and re-seeded into 96-well plates. The appropriate peptide ligands were then applied at a final concentration range of 1 nM to 1 mM. The cells were cultured for an additional 12 h to allow for the induction and expression of luciferase. Following incubation, the Bright-Glo Luciferase Assay System (Promega) was added to a final concentration of 5 μM. Luminescence was quantified using a VICTOR X5 multi-label plate reader (PerkinElmer) to assess activation of the sensors compared to their wild-type receptor counterparts.

### pH sensitivity

To evaluate the pH sensitivity of the neuropeptide sensors, HEK293T cells (either stably expressing the sensor or transiently transfected) were seeded into CellCarrier Ultra black-walled 96-well plates. EGFP^50^ and Gamillus^51^, each fused with a C-terminal CAAX motif to ensure plasma membrane localization (EGFP-CAAX and Gamillus-CAAX, respectively), were utilized as control markers. A series of Tyrode’s or PBS buffers was prepared with pH values ranging from 2.0 to 9.0 (2.0, 2.5, 3.0, 3.5, 4.0, 4.5, 5.0, 6.0, 7.0, 8.0, and 9.0). To equilibrate the intracellular pH with the extracellular environment, each buffer was supplemented with 15 μM (final concentration) digitonin to permeabilize the plasma membrane. Fluorescence spectra were recorded using a Safire2 multi-mode plate reader (Tecan). Imaging was performed using the Opera Phenix Plus or Operetta CLS high-content screening system. Prior to imaging, the culture medium was replaced with the digitonin-containing pH buffers, and the cells were incubated for 5 minutes. Each pH condition was measured in at least three replicate wells. Fluorescence images were first acquired and analyzed in the absence of ligand, followed by measurements in the presence of a saturating concentration of the corresponding neuropeptide ligand.

### Fiber photometry recording

*In vivo* fluorescence signals were recorded using a multi-channel fiber photometry system (Inper, Hangzhou, China) equipped with high-sensitivity, low-noise CMOS sensors. To enable excitation-ratiometric imaging of the neuropeptide sensors, two independent LED sources—488 nm and 405 nm—were used to excite the deprotonated and protonated states of the fluorescent reporter, respectively. The system was configured with a sampling frequency of 10 Hz and supported multi-site or multi-region recording via bundled optical fibers.

Raw photometry data were processed using the InperDataAnalysis software and Spike 2. For both the 488-nm and 405-nm channels, the signals were subjected to background subtraction and bi-exponential fitting to correct for photobleaching. The ratiometric signal (R) was calculated as the ratio of the 488-nm-excited fluorescence to the 405-nm-excited fluorescence (F488/F405), providing a self-calibrating measurement resistant to motion artifacts and expression differences. For quantitative analysis, the change in fluorescence was expressed as ΔF/F_0_ or ΔR/R_0_, where F_0_ and R_0_ represent the baseline values calculated from the mean signal during a stable period preceding each event.

### Tail suspension test

To evaluate the sensors’ change in fluorescence in response to acute stress, the mice were subjected to a tail suspension test. Prior to recording, the mice underwent a 2-day acclimation period as follows: on day 1, the mice were tethered to the optical fiber and allowed to explore a neutral environment for 30 min; on day 2, the mice were habituated to the experimental behavioral chamber for 30 min. During the test, the researcher suspended the mouse by the tail at a height of 60 cm for 1 min. Each mouse underwent 5–6 trials per session with a 5 min inter-trial interval. A 15-min baseline was recorded prior to the first trial, and recording continued for 15 min after the final trial. The baseline (F_0_ or R_0_) for each trial was defined as the average signal during the 1 min preceding the suspension event.

### Isoflurane anesthesia

To measure neuropeptide dynamics during state transitions, the mice were subjected to isoflurane-induced anesthesia using a multi-channel system (RWD Life Science). After a 1-hour baseline recording in the behavioral chamber, anesthesia was induced with 1.5% isoflurane delivered at an oxygen flow rate of 0.7 L/min for 2 h. Following the anesthesia period, recording continued for 3 h to capture the recovery phase until the mice returned to normal activity levels. For long-term data analysis, the raw signals were exported as CSV format and processed in Spike 2. To facilitate visualization of hour-scale dynamics, the data were decimated by a factor of 10 and smoothed using a 1-s moving average.

### Psychoactive drug administration

To assess sensor performance during drug-induced behavioral states, the mice received intraperitoneal injections of cocaine (20 mg/kg body weight); control groups received a 300 ml injection of sterile saline. For these experiments, a 20-min baseline was recorded, followed by 60 min of post-injection recording. The experimental design of repeated cocaine injections was as follows: the mice received a saline injection on Day 1, followed by injections of 20 mg/kg cocaine on Day 2 and Day 8. Simultaneously, locomotor activity was quantified using the ezTrack tool, with total distance traveled and animal trajectories calculated in 5-min bins to correlate neuropeptide dynamics with drug-induced behavioral changes.

### Quantification and statistical analysis

Experimental groups for both the *in vitro* and *in vivo* studies were assigned randomly. Sample sizes were determined based on established protocols from previous reports (Sun et al., 2020). Fluorescence imaging data obtained from HEK293T cells and cultured primary neurons were processed using ImageJ (NIH) and further analyzed using custom-written MATLAB scripts (MathWorks).

For the ratiometric analysis, the fluorescence signals from the two excitation wavelengths were baseline-subtracted and used to calculate the ratio R (F488/F405). To account for slight photobleaching, the signals were corrected using an exponential-function fit. Background subtraction was performed by measuring non-fluorescent regions adjacent to the regions of interest (ROIs) using ImageJ.

Statistical analyses were performed using Origin 2020 (OriginLab) and Prism 8/9 (GraphPad). Unless specified otherwise, all summary data are expressed as the mean and SEM. For comparisons between two groups, the paired or unpaired Student’s *t*-test was. Comparisons among multiple groups were performed using a one-way analysis of variance (ANOVA). All statistical tests were two-tailed, and differences with a *P*-value <0.05 were considered significant.

---

**Fig. S1.**
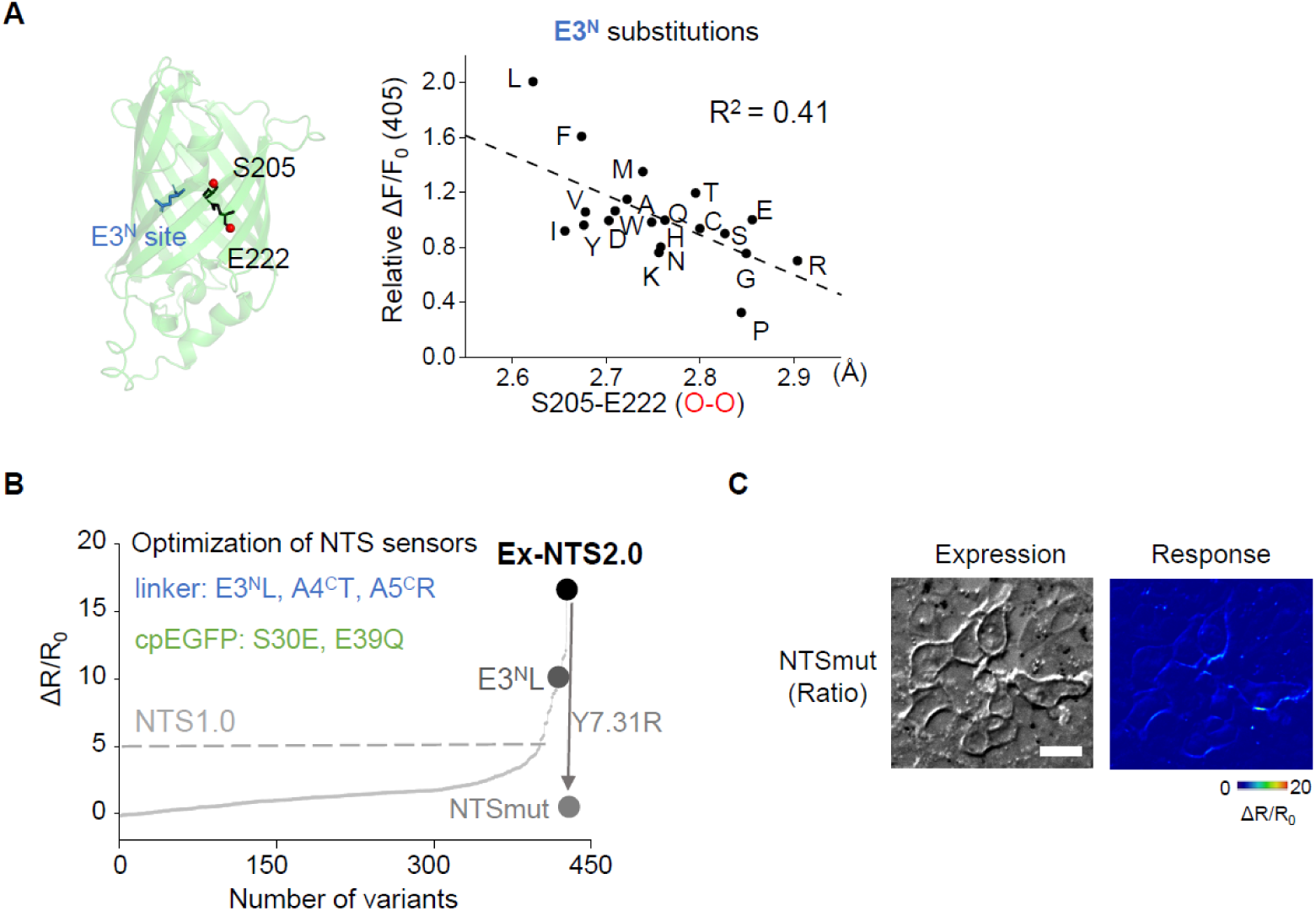
Characterization of the virtual screening pipeline and generation of Ex-NTS2.0 variants A. Structural modeling and correlation analysis for the E3^N^ site. Left: Structure of the cpEGFP part of the GRAB sensor predicted by Alphafold3 highlighting the E3^N^ site on the cpEGFP scaffold. Right: Correlation between the predicted S205–E222 atom distance (O–O, in Å) from virtual screening and the experimental fluorescence response (ΔF/F_0_) under 405 nm excitation for E3^N^ variants. (B) Summary of NTS sensor optimization. The non-responsive control, NTSmut, was generated by introducing the Y7.31R mutation. (C) Characterization of the NTSmut control. Representative images showing cell surface expression (left) and the lack of fluorescence response (ΔR/R_0_) of NTSmut in HEK293T cells. Scale bar, 20 μm.

**Fig. S2.**
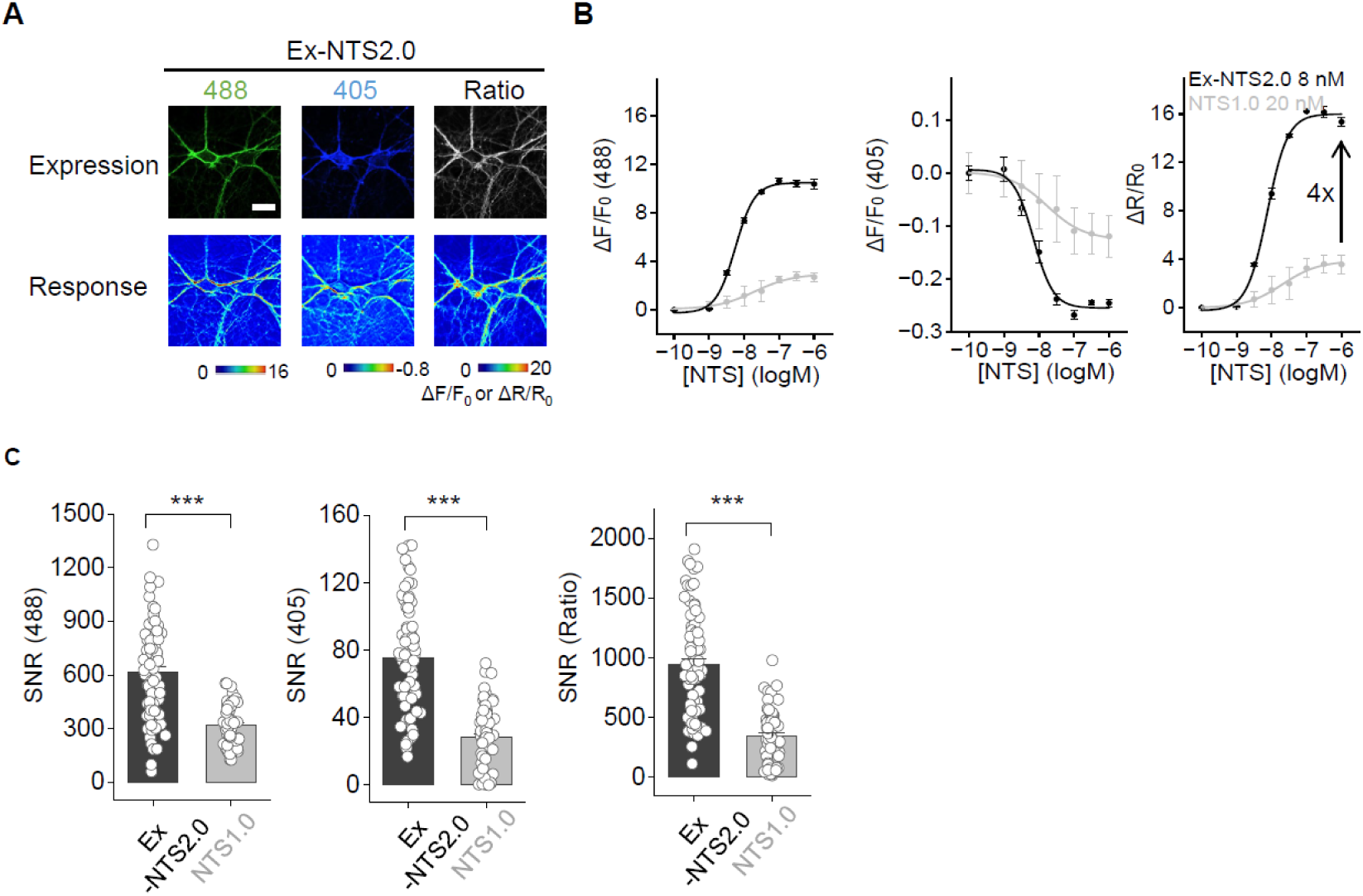
Characterization of the Ex-NTS2.0 sensor in cultured neurons A. Expression and fluorescence response of Ex-NTS2.0. Confocal images showing the expression (top) and corresponding NTS-induced fluorescence responses (bottom, ΔF/F_0_ or ΔR/R_0_) of Ex-NTS2.0 in cultured neurons across the 488 nm channel, 405 nm channel, and ratiometric signal. Scale bar, 20 μm. B. Dose-response curves of Ex-NTS2.0 and NTS1.0. Concentration-dependent fluorescence changes (ΔF/F_0_ for 488 nm and 405 nm; ΔR/R_0_ for Ratio) plotted against NTS concentrations for Ex-NTS2.0 (black) and NTS1.0 (gray). C. Comparison of Signal-to-Noise Ratio (SNR). Quantitative summary of the SNR between Ex-NTS2.0 (dark gray) and NTS1.0 (light gray) in the 488 nm, 405 nm, and ratiometric channels. Individual data points represent single ROI measurements. n = 3 coverslips per group. \*\*\**P*<0.001 (two-tailed Student’s *t*-test)

**Fig. S3.**
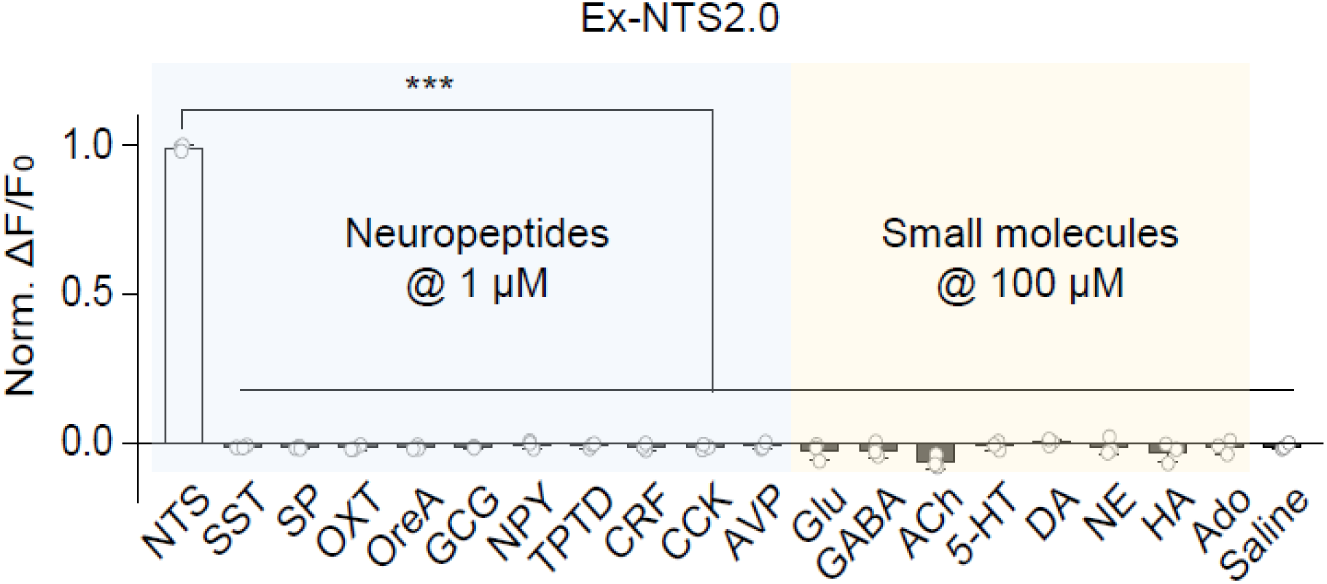
Specificity of Ex-NTS2.0 Screening of Ex-NTS2.0 against various neurochemicals. Normalized fluorescence response of Ex-NTS2.0-expressing cells treated with various potential competitors. Left panel (shaded light blue) shows responses to a panel of non-target neuropeptides (SST, SP, OXT, OxA, GCG, NPY, TPTD, CRF, CCK, AVP) applied at 1 μM. Right panel (shaded light yellow) shows responses to common small-molecule neurotransmitters and modulators (Glu, GABA, ACh, 5-HT, DA, NE, HA, Ado) applied at 100 μM, with saline serving as a vehicle control. \*\*\**P*<0.001 (one-way repeated measures ANOVA followed by Tukey’s multiple-comparison tests).

**Fig. S4.**
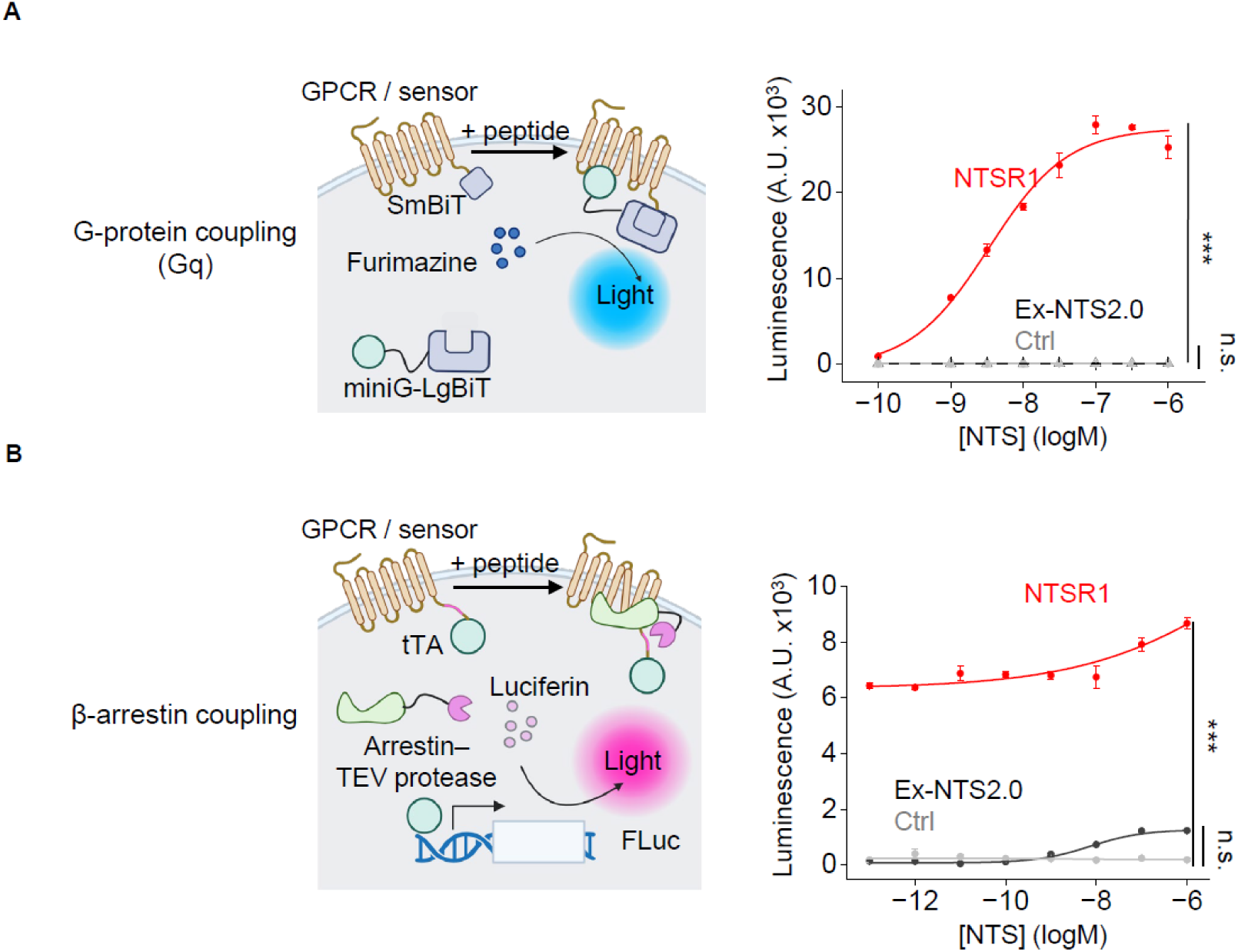
Downstream signaling and G-protein / β-arrestin coupling tests A. G-protein coupling assay. Left: Schematic diagram of the split-luciferase complementation assay (NanoBiT) used to measure G-protein recruitment. Right: Dose-response curves of luminescence for NTSR1 (red), Ex-NTS2.0 (black), and control (Ctrl, gray). \*\*\**P*<0.001, and n.s., not significant. B. β-arrestin coupling assay. Left: Schematic representation of the Tango assay utilized to monitor β-arrestin recruitment. Right: Luminescence dose-response curves following NTS application for NTSR1 (red), Ex-NTS2.0 (black), and control (Ctrl, gray) groups. \*\*\**P*<0.001, and n.s., not significant.

**Fig. S5.**
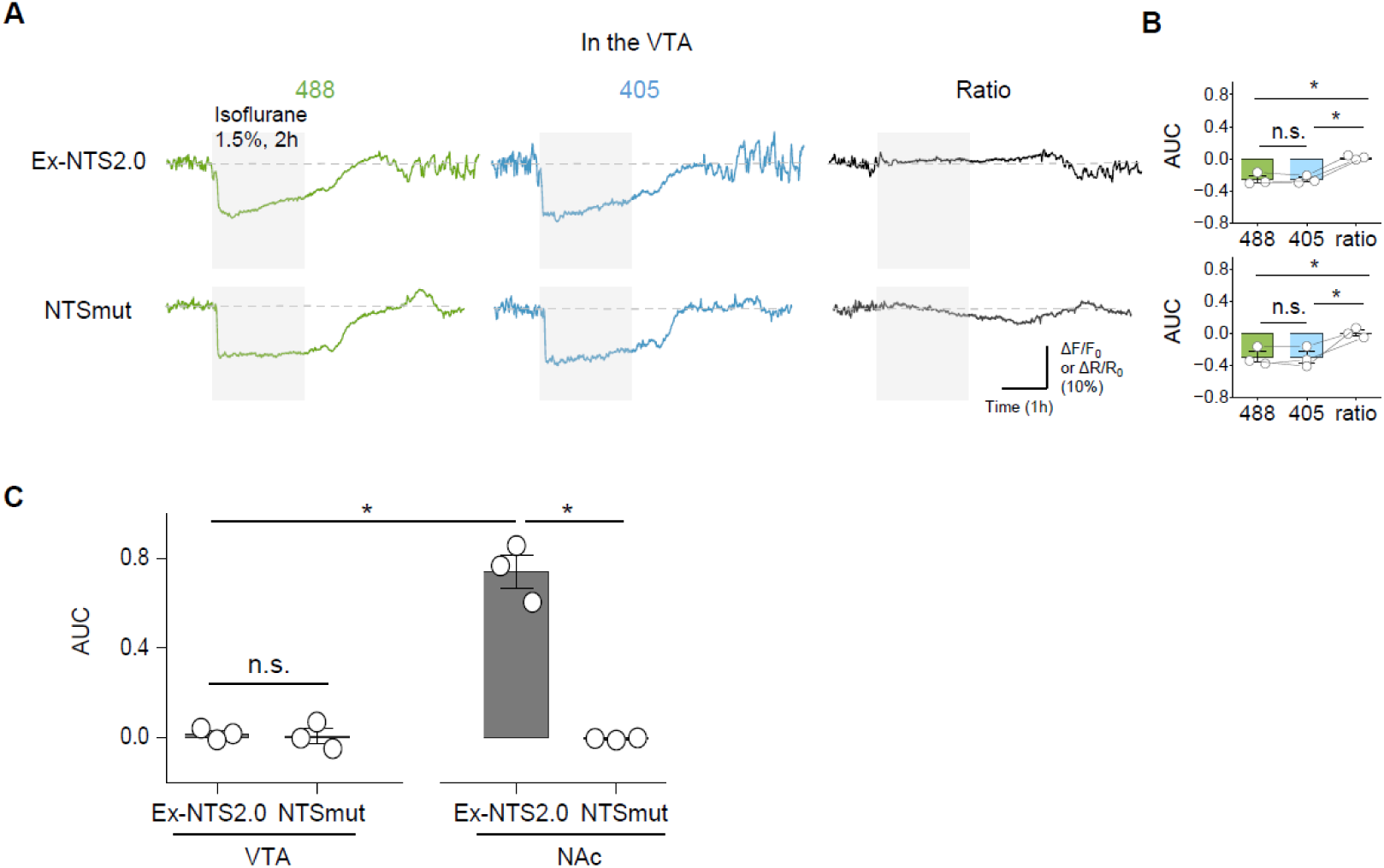
Ratiometric recording of Ex-NTS2.0 and NTSmut in the VTA during anesthesia A. Representative fluorescence and ratiometric traces during isoflurane-induced anesthesia. In vivo fiber photometry traces of Ex-NTS2.0 (top) and NTSmut (bottom) expressed in the VTA. Shaded gray bars indicate the duration of isoflurane administration (1.5%, 2 h). Traces display fluorescence changes under 488 nm (green) and 405 nm (blue) excitation, alongside the calculated ratiometric signal (Ratio, black). B. Quantification of fluorescence channel responses in the VTA. Summary data showing the area under the curve (AUC) during the anesthesia period for the 488 nm, 405 nm, and ratiometric signals of Ex-NTS2.0 (top) and NTSmut (bottom). C. Comparison of ratiometric AUC responses between VTA and NAc. Group data quantifying the ratiometric AUC for Ex-NTS2.0 and NTSmut in the VTA versus the NAc during anesthesia. The data in the NAc were replotted from Fig. 3D. Individual data points represent single animals. \**P*<0.05 and n.s., not significant.

**Fig. S6.**
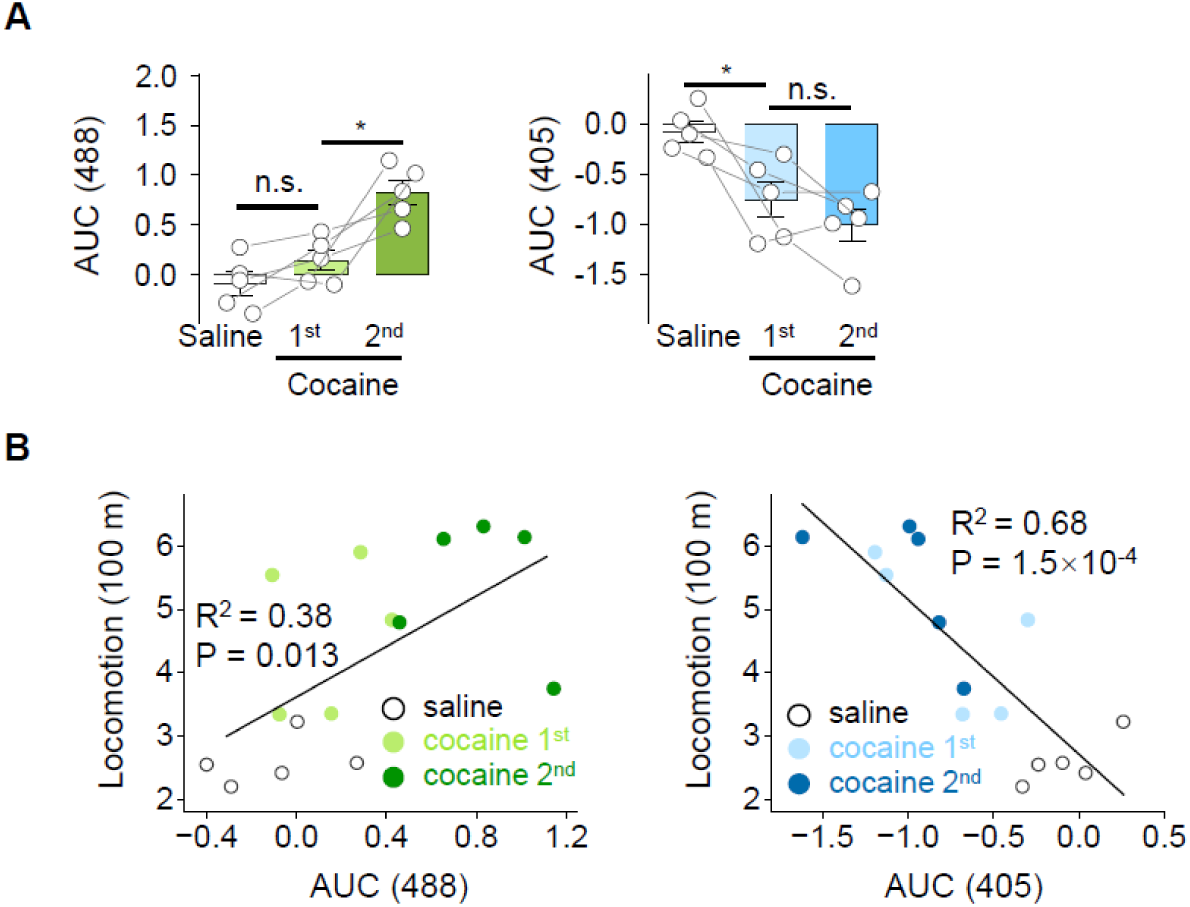
Fluorescence signals and locomotion tracking under cocaine treatment A. Quantification of Ex-NTS2.0 fluorescence responses via AUC analysis. Area under the curve (AUC) measurements for the 488 nm channel (left, green) and 405 nm channel (right, blue) comparing saline, 1^st^ cocaine, and 2^nd^ cocaine administrations. Connected lines represent individual animals. \**P*<0.05 and n.s., not significant. B. Correlation analysis between fluorescence signals and locomotion. Linear regression plots correlating total locomotion distance with the AUC of the 488 nm channel (left) and the AUC of the 405 nm channel (right). Color-coded symbols distinguish saline control (open shapes), 1st cocaine (light color), and 2nd cocaine (dark color) conditions across trials.

**Fig. S7.**
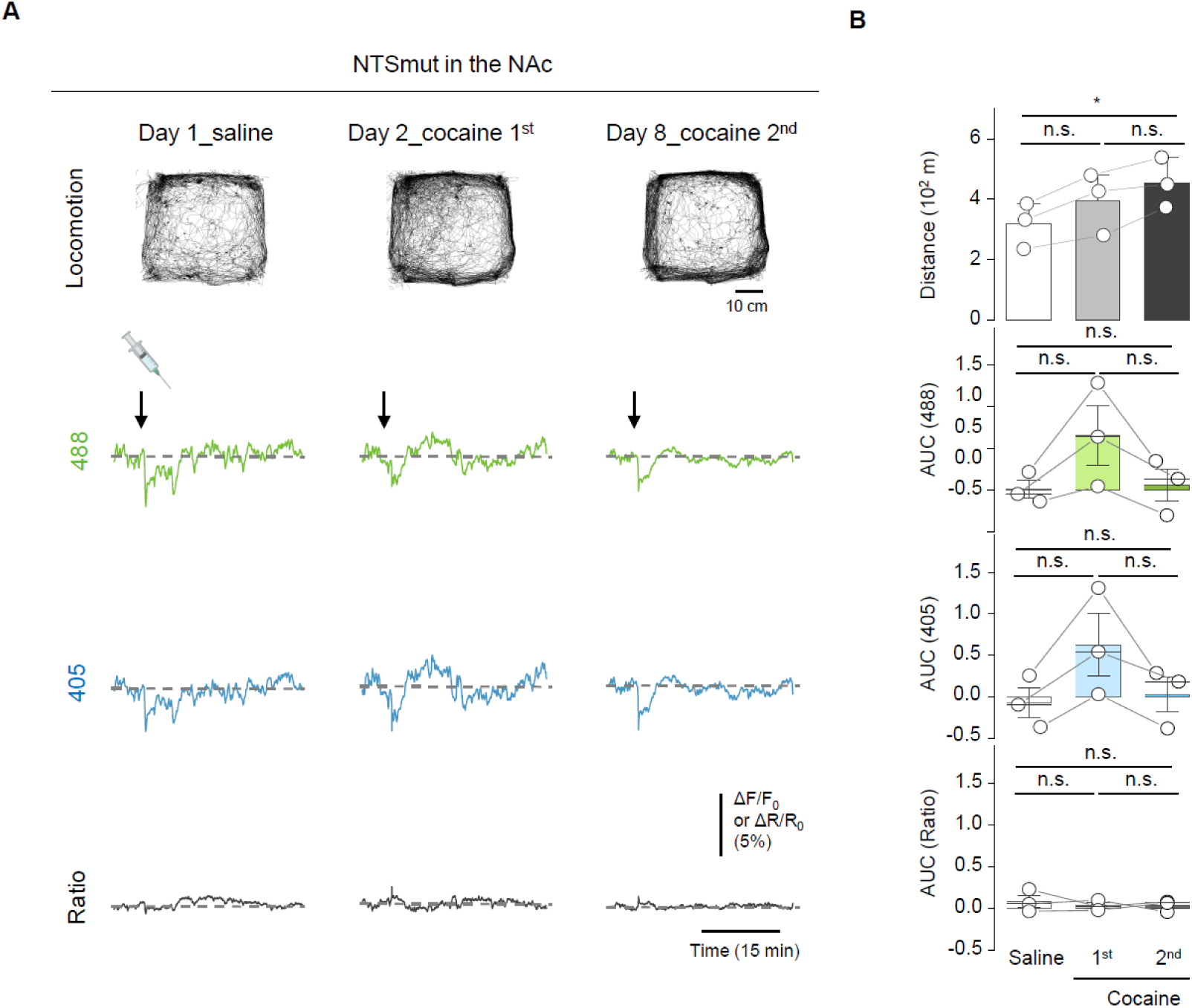
NTSmut exhibits no significant response during cocaine administration A. Locomotion tracks and NTSmut fluorescence traces in the NAc. Top: Open-field locomotion tracks across three sessions: Day 1 (saline), Day 2 (1st cocaine), and Day 8 (2nd cocaine). Scale bar, 10 cm. Bottom: Simultaneous in vivo fiber photometry traces of NTSmut in the NAc. Arrows denote injection time. Traces show responses under 488 nm (green), 405 nm (blue) excitation, and the ratiometric signal (Ratio, black). B. Quantification of behavioral and photometric responses. Statistical summaries comparing saline, 1st cocaine, and 2nd cocaine sessions. Top: Total locomotion distance. Bottom three panels: Area under the curve (AUC) quantification for the 488 nm channel, 405 nm channel, and ratiometric signal. Circles represent individual animals. \**P*<0.05 and n.s., not significant

## Notes

### Competing Interest Statement

The authors have declared no competing interest.

